# Removing the hidden data dependency of DIA with predicted spectral libraries

**DOI:** 10.1101/681429

**Authors:** B. Van Puyvelde, S. Willems, R. Gabriels, S. Daled, L. De Clerck, S. Vande Casteele, A. Staes, F. Impens, D. Deforce, L. Martens, S. Degroeve, M. Dhaenens

**Affiliations:** ProGenTomics, Laboratory of Pharmaceutical Biotechnology, Ghent University, Ghent, Belgium; VIB-UGent Center for Medical Biotechnology, Ghent, Belgium; Department of Biomolecular Medicine, Ghent University, Ghent, Belgium; VIB Proteomics Core, Ghent, Belgium

**Author notes:** Authors contributed equally (B. Van Puyvelde, S. Willems, R. Gabriels).

## Abstract

Data-Independent Acquisition (DIA) generates comprehensive yet complex mass spectrometric data, which imposes the use of data-dependent acquisition (DDA) libraries for deep peptide-centric detection. We here show that DIA can be redeemed from this dependency by combining predicted fragment intensities and retention times with narrow window DIA. This eliminates variation in library building and omits stochastic sampling, finally making the DIA workflow fully deterministic. Especially for clinical proteomics, this has the potential to facilitate inter-laboratory comparison.

**Significance of the Study:** Data-independent acquisition (DIA) is quickly developing into the most comprehensive strategy to analyse a sample on a mass spectrometer. Correspondingly, a wave of data analysis strategies has followed suit, improving the yield from DIA experiments with each iteration. As a result, a worldwide wave of investments in DIA is already taking place in anticipation of clinical applications. Yet, there is considerable confusion about the most useful and efficient way to handle DIA data, given the plethora of possible approaches with little regard for compatibility and complementarity. In our manuscript, we outline the currently available peptide-centric DIA data analysis strategies in a unified graphic called the DIAmond DIAgram. This leads us to an innovative and easily adoptable approach based on predicted spectral information. Most importantly, our contribution removes what is arguably the biggest bottleneck in the field: the current need for Data Dependent Acquisition (DDA) prior to DIA analysis. Fractionation, stochastic data acquisition, processing and identification all introduce bias in the library. By generating libraries through data independent, i.e. deterministic acquisition, stochastic sampling in the DIA workflow is now fully omitted. This is a crucial step towards increased standardization. Additionally, our results demonstrate that a proteome-wide predicted spectral library can surrogate an exhaustive DDA Pan-Human library that was built based on 331 prior DDA runs.

## Article

With DIA, an MS instrument regularly measures precursor ions and continuously cycles through predefined mass over charge ratio (*m/z*) windows to equally regularly measure the intensity of their fragment ions throughout a liquid chromatography (LC) gradient. This is both more qualitative and quantitative than data-dependent acquisition (DDA), where precursor ions are measured intermittently while fragment ions are only measured stochastically. However, the complexity of DIA data has shown to be very challenging.

To date, the most common way to address this complexity is using previously identified peptides from DDA as targets in the DIA data. First, DDA peptide identifications are translated into a spectral library with Peptide Query Parameters (PQPs), which typically contain the sequence as well as the analytical coordinates (*m/z*, intensity, and retention time or RT) for the observed ions for a given peptide. These PQPs are then used to compute an evidence score for each target peptide, based on its fragment traces in DIA ^[1]^. Ultimately, these evidence scores are supplemented with additional features, e.g. ppm and RT errors, allowing a semi-supervised machine learning algorithm to weigh and re-score the target peptides to obtain a maximum of true targets at an empirically determined False Discovery Rate (FDR) using the target-decoy approach ^[2][3][4]^.

Unfortunately, deriving PQPs from DDA data intrinsically means transferring its limitations. In fact, fractionation, stochastic data acquisition, processing and identification introduce bias in the library and require considerable effort. This compromises inter-laboratory comparison and can even alter the biological conclusions between labs ^[5]^. However, thanks to the availability of state-of-the-art prediction algorithms, these PQPs can now be predicted directly, setting the stage for much easier and much more reproducible peptide-centric DIA data extraction ^[6][7][8]^.

Here, we compare the effect of using libraries from different origins on peptide-centric approaches, by assessing their qualitative and quantitative performance on a public wide window (10 - 20 *m/z*) DIA dataset of HeLA cells ^[9]^ (Figure 1). Three basic spectral libraries were used here, with PQPs derived from (a) an experimental DDA dataset, (b) a protein sequence database (FASTA), and (c) a predicted spectral dataset. Each of these three libraries can be used directly as a source library, or can be converted into a DIA library by using them first on a narrow window (2 *m/*z) DIA dataset of the sample. The resulting six possible libraries can all be used alike by the EncyclopeDIA software to identify and quantify wide window DIA data ^[9]^.

**Figure 1.**
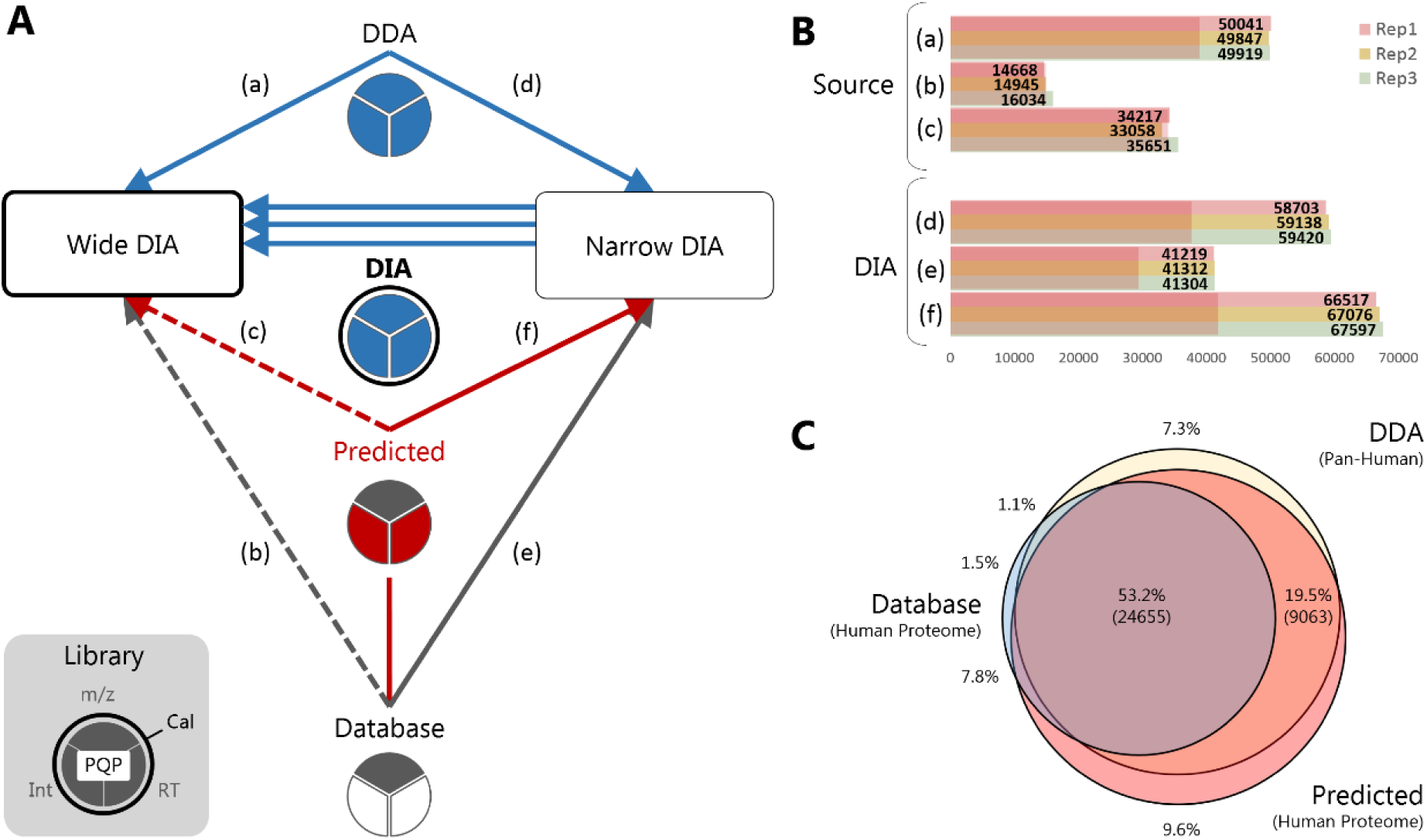
Peptide-centric data extraction from wide window DIA data. (A) DIAmond DIAgram presenting peptide-centric strategies for DIA data extraction. Peptide-centric approaches rely on libraries **(central column)** that contain Peptide Query Parameters (PQPs) which are derived from the peptide sequence and can additionally contain the three ion coordinates, i.e. mass to charge ratio **(*m/z*)**, Intensity **(Int)** and retention time **(RT) (three-part pie charts)**. These can either be experimental **(blue)**, theoretical **(grey)**, or predicted **(red)**. PQPs are used to score the evidence of peptide detections in continuous DIA data **(boxes)**. These are supplemented with additional features of the match so that a support vector machine can weigh and re-score them to obtain a maximum of true targets at an empirically determined FDR using the target-decoy approach **(arrow heads). DDA** source libraries (both in-house and public) only comprise prior proteotypic peptide identifications and contain measured PQPs for all three ion coordinates. These are therefore directly applicable to quantify peptides in 10 – 20 *m/z* wide window DIA **(Wide DIA)** data **(a)**. However, when a proteome **FASTA** is used as a source library, sensitivity is reduced **(dashed arrow)**, i.e. too many false negatives are produced due to the high statistical burden **(b)**. This also holds for libraries with **predicted** fragment intensities (MS²PIP) and RT (Elude), albeit to a lesser extent **(c)**. Prior 2 *m/z* narrow window DIA **(Narrow DIA)** provides the specificity to remove false targets in the sample first **(d)(e)(f)**. The DIA ion coordinates from these detections can additionally be integrated into new and calibrated PQPs **(cal)**. These **DIA** libraries, called chromatogram libraries, can be derived from any source library **(triple arrow). (B)** Doubly and triply charged peptide detections in wide window DIA following each of the routes depicted in **(A)**. Shading highlights the number of peptides that is detected in triplicate wide window DIA runs with at least three transitions, allowing robust quantification. **(C)** Comparison of the identified peptide sequences in Wide DIA for route **(d), (e)** and **(f)**. The large overlap shows that all three approaches detect proteotypic peptides. Only peptides of double and triple charge that are detected in triplicate wide window DIA runs with at least three transitions are shown.

In-house or public DDA source libraries are frequently built by extensive fractionation of samples. With adequate statistical control, such proteotypic libraries allow direct peptide detections in wide window DIA (Figure 1Aa) ^[10]^. We illustrate this by using the publically available Pan-Human library, which contains nearly 10.000 proteins derived from 331 DDA runs on a range of human cell lines and tissues ^[11]^ (Figure 1Ba). To reduce the effort and variability from DDA library building, a library-free peptide-centric data analysis workflow was proposed recently ^[12]^. Herein, the PECAN (or Walnut) scoring algorithm allows direct detection of peptides derived from a FASTA in wide window DIA data (Figure 1Ab). This is akin to a source library that (i) contains only peptide sequences and *m/z* coordinates, and (ii) lacks prior selection of proteotypic peptides. On wide window DIA data this approach thus provides a limited number of PQPs, which is not sufficient to differentiate between the high number of false targets, i.e. true negatives, and the lower number of true positives in the library ^[13]^. This manifests as indiscernible target and decoy score distributions, resulting in a very high False Negative Rate (FNR) (Figure 1Bb).

Here we propose a promising way to improve upon the FASTA source library - while still omitting prior DDA - by predicting fragment ion intensity and RT *in silico* (Figure 1Ac and Figures S1–S2). Using a spectral dataset with such predicted fragment intensities (MS²PIP) and peptide RTs (Elude) more than doubles the number of peptides detected in the wide window DIA (Figure 1Bc) ^[6][14]^. However, considering all tryptic peptides in a Human proteome still underperforms compared to the Pan-Human DDA library, which is fully contained in the predicted spectral dataset (Figure 1Ba and 1Bc). Notably, this is not due to poor prediction because predicting only those peptides present in the Pan-Human library performs very similar to using the Pan-Human library directly (Figure S3) and the underperformance can thus only be attributed to the many false targets when using the complete database ^[10]^. An elegant way to filter out false target peptides upfront, is by measuring a pool from every condition with staggered narrow window DIA (Figure 1Ad, 1Ae and 1Af). This reduces MS2 chimericity to DDA-like quality in a DIA setting, allowing detection with increased specificity. This accurate prior filtering makes the statistical burden of false targets in the wide window DIA surmountable again. Notably, due to instrument limitations this *Precursor Acquisition Independent From Ion Count* (PAcIFIC) ^[15]^ can currently only be performed by means of gas phase fractionation (GPF), i.e. sampling different *m/z* regions separately ^[9]^. Still, the added acquisition depth and specificity allows the detection of 88k (DDA), 47k (FASTA) and 95k (predicted) peptides in six narrow window GPF DIA runs of a HeLA cell lysate (Figure S4). To assure that this additional filtering is accurate, we confirmed the estimated FDR by using an entrapment experiment wherein we included *Pyrococcus furiosus* proteins as false targets alongside the expected human proteins in the respective source libraries ^[16]^. Hereby, the measured FDR for narrow window DIA filtering is 2% for the DDA, 1% for the FASTA, and 1% for the predicted source library, in accordance with the theoretically estimated FDR based on the target-decoy strategy. In the process, we can measure the identification cost of adding false targets: adding 3-6% false targets results in an average decrease of 1-2% in detections (see Entrapment section in Methods).

Additionally, the peptide detections in narrow window DIA can be translated into novel and integrated PQPs, which are calibrated to the specific LCMS system and are specific to DIA (Figure 1A). This approach was recently made readily applicable as chromatogram libraries: DIA libraries of narrow window DIA peptide detections comprising their calibrated PQPs ^[9]^. Such chromatogram libraries outperform direct wide window DIA extraction for every source library. The modest gain for a DDA source library (~20%) derives mainly from PQP calibration, as only 50% of the source peptides was filtered out (Figure 1Ba and 1Bd). In contrast, in the FASTA source library, 98,5% of the peptides were filtered out, and RT and intensity coordinates were generated *de novo*. Taken together, this resulted in the largest gain (~170%) (Figure 1Bb and 1Be). Finally, the chromatogram library derived from a predicted spectral library increases the number of detections by ~100% compared to direct wide window DIA data extraction, making it the most efficient overall peptide detection strategy of the DIAmond DIAgram (Figure 1Bc and 1Bf). The large overlap between the peptide sequences detected by all three chromatogram libraries convincingly shows that the Pan-Human library is very exhaustive and that all three chromatogram libraries mainly detect proteotypic peptides (Figure 1C). Peptides unique to the Pan-Human library include very high molecular masses that were not predicted, high molecular weight peptides that generate many doubly charged transitions that are not predicted by default, as well as very small peptides with inherently poor RT or fragmentation pattern predictions. Peptides that are unique to the predicted library are mainly peptides not present in the Pan-Human source library. Importantly, the PQP requirements of the source library for building chromatogram libraries on narrow window DIA are relatively liberal: the measured Pan-Human library was acquired on a TripleTOF instrument but allows wide window DIA data peptide detection on an Orbitrap instrument. The *in silico* equivalent is that 95% of the detected peptides overlap when the MS²PIP engine is trained on either Orbitrap or TripleTOF data. As a result, other fragment ion intensity predictors such as Prosit and Deep Mass ^[7][8]^ perform similarly when combined with narrow window DIA ^[17]^ (Figure S5).

We therefore conclude that predicted libraries are highly relevant and performant for wide window DIA identification, and that three elements of a spectral library affect its overall performance: (i) the amount of false targets included, (ii) the amount of informative PQPs, and (iii) the accuracy of PQPs on the specific instrument setup. In this study, we could show that a narrow window DIA acquisition of six GPFs combined with a predicted spectral library of the full human proteome was able to surrogate a measured DDA Pan-Human library, thus liberating the DIA workflow from any stochastic acquisition. Especially for clinical proteomics, this can facilitate inter-laboratory comparison. Importantly, the software tools MS²PIP, ELUDE and EncyclopeDIA are all instrument independent, publicly available, and mutually compatible, thus making this workflow immediately accessible to everybody interested.

## Code availability

MS²PIP, Elude and EncyclopeDIA are open source, licensed under the Apache-2.0 License, and are hosted on https://github.com/compomics/ms2pip_c, https://github.com/percolator/percolator and https://bitbucket.org/searleb/encyclopedia/wiki/Home. All supporting material is available on https://github.com/brvpuyve/MS2PIP-for-DIA/.

## Acknowledgments

This research was mainly funded by mandates from the Research Foundation Flanders (FWO) awarded to BVP [grant number 11B4518N], RG [grant number 1S50918N] and MD [12E9716N]. Partial funding was received through project grants from the FWO [G013916N and G042518N], from the European Union’s Horizon 2020 Program under Grant Agreement 823839 [H2020-INFRAIA-2018-1], and from a PhD grant from the Flanders Agency Entrepreneurship and Innovation (VLAIO) awarded to LDC [SB-141209].

## Competing interests

The authors have declared no conflict of interest.

## Author Contributions

BVP performed all data analysis at the ProGenTomics facilities. The initial experimental design was conceived and performed by BVP, SW, MD, SDa, LDC, AS, DD and FI. RG, SDe and LM performed all machine learning predictions. MD, BVP, RG and SW wrote the draft manuscript. All authors provided critical feedback during research and writing. MD conceived the idea of using predicted libraries for DIA data extraction and supervised the project.

## SUPPORTING INFORMATION

### Introduction

DIA data has been presented as a permanent record of everything. Thus, applying our novel approach can significantly broaden the biological perspective on newly acquired as well as existing data. Using predicted spectral libraries to replace measured DDA libraries not only reduces workload and increases reproducibility; it will also facilitate the implementation of DIA into more applied fields such as clinical proteomics. Since the software tools MS²PIP, Elude and EncyclopeDIA are instrument independent, publicly available and mutually compatible, the presented workflow is accessible to everybody and directly applicable ^[1–3]^. Therefore, we present this methods section in the form of a systematic tutorial. Briefly, both source and DIA libraries can be used in EncyclopeDIA to detect peptides in wide window DIA. However, converting source libraries into a DIA library will significantly improve the number of peptides that can be detected. This requires an additional narrow window DIA of several gas phase fractions (GPF) of a mixture of the samples. When these GPFs are acquired in the same batch as the wide window DIA, the benefit of PQP calibration is maximized.

All external resources are available on GitHub https://github.com/brvpuyve/MS2PIP-for-DIA for reproducibility.

### Prediction Models: Elude and MS²PIP

#### Elude: Retention Time prediction

For RT prediction, we employed Elude (version 3.02), which is available from the Percolator GitHub repository (https://github.com/percolator/percolator/releases) ^[2]^.

We trained an Elude model on the Pan-Human spectral library ^[4]^. The spectral library was downloaded from SWATHAtlas in SpectraST SPTXT file format. The peptide sequences and their respective RTs were parsed from the SPTXT file to an MS²PIP PEPREC file using the speclib_to_mgf.py script, which is available in the conversion_tools folder of the MS²PIP GitHub repository. Out of all consensus peptide spectra built from five or more identified spectra, 10000 peptides and their mean RTs were randomly sampled for training, 10000 were randomly sampled for testing and all remaining were used for final validation of the model. The training, test and validation datasets were converted and written to the Elude input file format. Through the Elude command line interface, we trained a model with the training and test subsets. Subsequently, we used the model to predict RTs for the validation subset of the dataset. The median absolute difference in experimental and predicted RTs (DeltaRT) of the validation dataset was 3.2 minutes and 95% of the DeltaRTs were less then 12.1 minutes (Figure S1). The model predictions have a Spearman rank correlation with the validation RTs of 0.98.

The spectral library contains oxidation and carbamidomethylation peptide modifications. As a result, the currently trained Elude model is only able to predict RTs for unmodified peptides and peptides containing these modifications. The RTs included in the original Pan-Human SPTXT spectral library are normalized to the iRT Kit peptide sequences by SpectraST. All RT values predicted by the Elude model therefore take over this normalization. As is the case for experimental RTs, the predicted RTs are aligned to the experimental dataset by EncyclopeDIA. The Elude model file is available on our GitHub repository.

#### MS²PIP: intensity Prediction

MS²PIP, the MS² Peak Intensity Predictor, first published by Degroeve et al., underwent significant improvements since its initial release in 2013 ^[5]^. Currently, a broad array of fragmentation models is available (e.g. Orbitrap-HCD, iontrap-CID, TripleTOF 5600+, …) ^[1]^. This gives the user the liberty to employ a model fit to the experimental setup. As both the narrow and wide window DIA datasets used in this project were obtained on a Q Exactive HF instrument (Thermo Fisher Scientific, Massachusetts, US), we employed MS²PIPs Orbitrap-HCD model, with the exception of the TT5600 model that was used for assessing PQP requirements (see main text). To further validate the application of this model, we calculated the correlations between MS²PIP predicted spectra and experimental spectra from the EncyclopeDIA DDA runs.

The Hela DDA dataset of the EncyclopeDIA article (MassIVE MSV000082805) was imported into Progenesis QI for Proteomics (Nonlinear Dynamics, Newcastle upon Tyne, UK) with default parameters. The peakpicked spectra were exported as .mgf and searched with Mascot 2.6.1 against the aforementioned human FASTA. Carbamidomethylation of Cysteine and oxidation of Methionine were respectively set as fixed and variable modifications. The precursor tolerance was set to 50 ppm and the fragment tolerance was set to 0.02 Da. The search included all 2+ and 3+ precursors, allowing up to 2 tryptic missed cleavages. Afterwards, the results were reimported into Progenesis QI for Proteomics and converted to an .msp spectral library.

The .msp spectral library was converted back to an .mgf and an MS²PIP PEPREC input format using the speclib_to_mgf.py script. Both files were then run through MS²PIP with the Orbitrap-HCD model, after which Pearson correlation coefficients (PCCs) were calculated for each experimental spectrum and its prediction. This resulted in a median PCC of 0.88 with an interquartile range of {0.795297, 0.938911} (Figure S2)

A second experiment was performed to evaluate the performance of predicted libraries. More specifically, as was done in Gessulat et al. ^[6]^, a clone of the Pan-Human library was produced using the HCD model and this was applied on the narrow-window DIA data, producing a chromatogram library containing 82.6k unique peptides. Afterwards, the Pan and Pan Clone chromatogram libraries were used in the peptide extraction of triplicate wide-window DIA runs. On average 63k and 62k peptides were identified at 1.0% FDR when searching the wide-window DIA data against the Pan-Human and the Pan Clone chromatogram library, respectively. The quantification reports on peptide and protein level were saved by EncyclopeDIA as .txt files and eventually imported in excel. Then, we manually filtered out all the peptide sequences with less than 3 fragment ions and those having an intensity of zero in at least one of the three replicates. The resulting reproducible peptide sequences were put in a Venn diagram to visualize the percentage overlap (Figure S3). The large overlap demonstrates i) the performance of the HCD fragmentation model of MS²PIP and ii) the retention time prediction of ELUDE to accurately mimic the fragmentation and retention time pattern of peptide sequences acquired on a TripleTOF instrument.

### Library Generation

#### DDA

An EncyclopeDIA .dlib version of the Pan-Human spectral library is publicly available on the EncyclopeDIA BitBucket homepage ^[4]^. This version contains 211k unique precursors (159k unique peptide sequences). Alternatively, EncyclopeDIA accepts Skyline .BLIB, Spectronaut .csv, MaxQuant msms.txt, .TraML and .msp files.

#### Database (FASTA)

Using a FASTA database does not require a separate library. More specifically, Walnut (a GUI re-implementation of the PECAN algorithm) is part of EncyclopeDIA and can directly detect peptides from DIA data using a FASTA database^[7]^.

Here, we used the human SwissProt proteome (UP000005640 downloaded on 12 February 2019, 20426 target sequences) downloaded as FASTA. The proteome was concatenated with the iRT FASTA obtained from the Biognosys webpage (on 12 February 2019) ^[8]^.

#### Predicted

Creating a predicted spectral library requires three steps: (i) creating an MS²PIP input PEPREC (peptide record) file from a FASTA, (ii) feeding that file to MS²PIP for predicting intensities and (iii) adding predicted retention times (RT) from Elude. For ease-of-use, we wrapped these three steps into a pipeline (fasta2speclib), that is included in the MS²PIP GitHub Repository.

MS²PIP is accessible either through the web server (https://iomics.ugent.be/ms2pip/) or via a local installation (https://github.com/compomics/MS2PIP_c/). A local installation is required to use the fasta2speclib pipeline. Here, MS²PIP (version 20190130) was downloaded and installed from the MS²PIP GitHub repository, as described in the extended install instructions. For RT prediction, we employed Elude version 3.02, which is available from the Percolator GitHub repository (https://github.com/percolator/percolator/releases).

Briefly, the fasta2speclib pipeline makes use of Biopython to read the FASTA and uses Pyteomics for the *in silico* digestion of the protein sequences ^[9,10]^. Next, redundant peptides and peptides not meeting the peptide length and precursor mass restrictions are removed from the peptide list. Following this step, all combinations of the requested charge states and modifications are added. Predicted spectra and RTs are then generated for all peptide-charge-modification combinations using MS²PIP and Elude. Finally, the results are written to a spectral library file (.msp, .mgf or .csv). Depending on the computational resources, a full human proteome can be predicted in just a few hours.

The fasta2speclib pipeline can be called through the command line interface as follows:

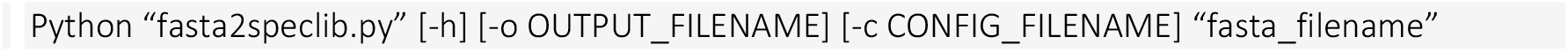

The results presented in this manuscript were generated by predicting a spectrum for every 2+ and 3+ tryptic peptide in the aforementioned FASTA, using the pre-trained MS²PIP Orbitrap-HCD model and the Elude RT. These models are described in more detail under “Prediction models”. Only tryptic peptides with a minimum length of 7 amino acid residues and a maximum precursor mass of 5000 Da were considered. Carbamidomethylation and oxidation were set as respectively fixed and variable modification, and two missed cleavages were allowed. The *in silico* spectral library was exported to an .msp file containing 3.3M precursors (between 400 – 1000 *m/z*). In the current version of MS²PIP (v20190624) the RT from Elude is automatically converted into minutes and written on a separate line in the .msp file. These predictions were performed on a Linux operated machine (Intel Xeon CPU X5670, 24 processors, 40 GB RAM) and took four hours.

#### DIA

DIA libraries, called chromatogram libraries, are generated by interrogating narrow window DIA data with any of the above source libraries. Details are described under “DIA data analysis: EncyclopeDIA”.

### RAW file processing

We used the publicly available dataset of the EncyclopeDIA article (MassIVE MSV000082805) of the HeLa S3 lysates to assay the different routes in the DIAmond DIAgram (Figure 1A, boxes). The three wide window DIA replicate runs were acquired with 25 overlapping 24 *m/z* windows and the staggered 4 *m/z* narrow window DIA data comprises six gas phase fractions (GPF) of 100 *m/z* each, together covering a 400 - 1000 *m/z* mass range. Following peak picking, these runs were demultiplexed into 12 *m/z* (wide DIA) and 2 *m/z* (narrow DIA) windows, respectively, and converted into mzML output files by MSConvertGUI with following parameters ^[11,12]^:

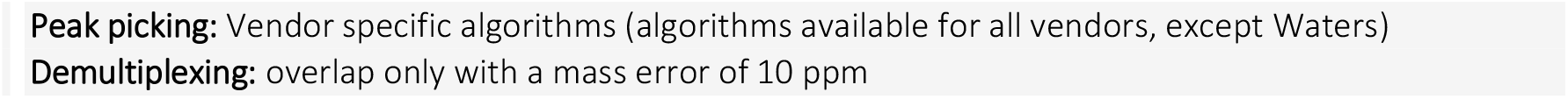

### DIA data analysis: EncyclopeDIA

We downloaded EncyclopeDIA from bitbucket (https://bitbucket.org/searleb/EncyclopeDIA/downloads/?tab=downloads) (version 0.8.2, 2019-05-21). EncyclopeDIA is a Java application developed to perform narrow- and wide window DIA data analysis. The application can be run on all three major operating systems (Windows, Mac and Linux), but in this project it was used on a Windows 7 operating system (Lenovo Thinkstation, Intel Xeon E5-2620 24 processors, 128 GB ram). EncyclopeDIA was operated through the graphical user interface but also comes with a command-line interface.

For applying EncyclopeDIA on predicted spectral libraries, the .msp file is first converted into a .dlib file using the conversion tool embedded in EncyclopeDIA. EncyclopeDIA also allows the conversion of other spectrum library formats into .dlib files.

General settings in EncyclopeDIA applied to all searches in this project are as follows:

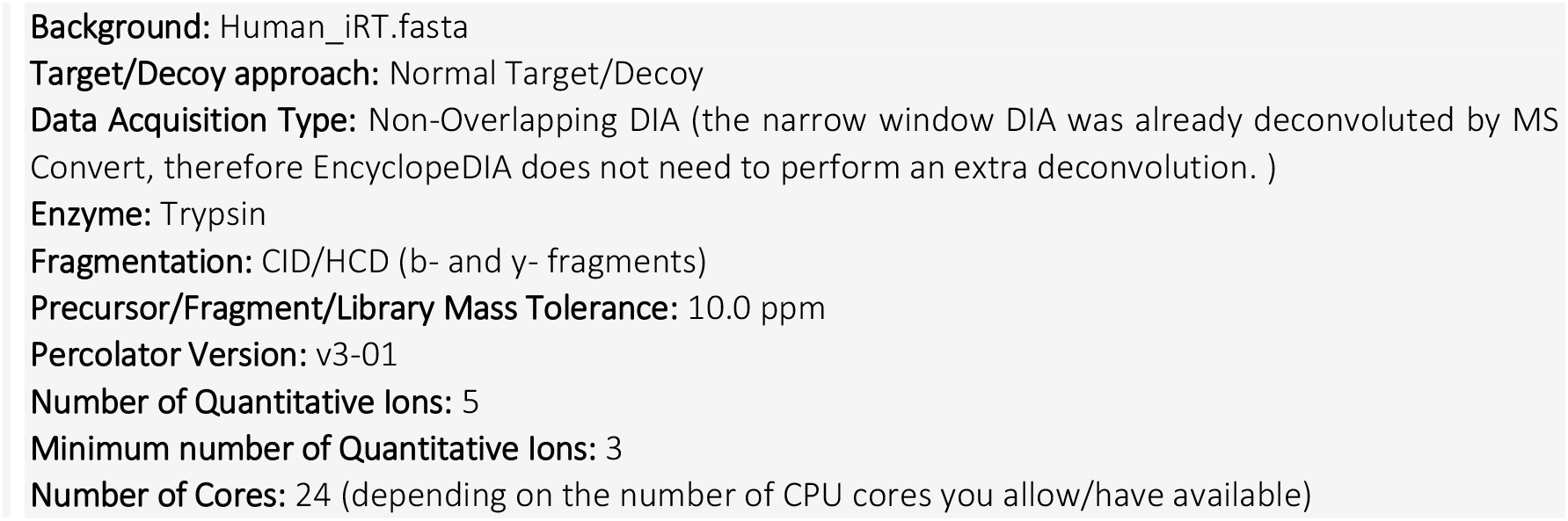

To allow direct comparison of all six routes of the DIAmond DIAgram, all libraries were trimmed upfront to retain only peptides in the 400 - 1000 *m/z* mass range. For the Pan-Human DDA library this results in 194k precursors, all charge states still included. Approximately 95% of the identified peptides on the wide window DIA were 2+ and 3+ and the other charge states were manually removed from the result file for comparison. The FASTA search was performed using Walnut, considering 2+ and 3+ precursors only. Finally, a third library was predicted by MS²PIP using the same FASTA. All three source libraries were separately used to detect peptides directly in the triplicate wide window HeLa DIA runs (Figure 1Aa-c). When the three source libraries were used to search the narrow window DIA data (Figure 1Ad, 1Ae, 1Af), this resulted in three DIA-based chromatogram libraries (.elib) of size 88k (DDA), 47k (FASTA) and the 95k (Predicted) peptides, respectively. In Figure S4, the overlap in peptide sequence is shown between the three chromatogram libraries. Subsequently, all three .elibs were used to search the wide window DIA data with the above parameters.

Figure 1B depicts the number of detected peptides in each replicate as reported by EncyclopeDIA. Additionally, the peptide quantification reports were exported as .txt files and peptide sequences with at least 3 transitions and non-zero intensities in all three wide window DIA samples were selected. These are represented as the shaded portion of the bar chart. Indeed, in most settings, only confident peptides that can be quantified with robust statistics and are detected in (almost) all runs, are useful. These recurring peptides equally have more robust FDR control. For this reason, we choose to focus only on these confident peptides in Figure 1C, as depicted in the figure caption. Note that the portion of unique peptides between robust detections in Pan-Human and predicted wide window DIA is considerably lower than in the chromatogram libraries that are intrinsically representing single detection. It would be interesting to investigate what the contribution of false detections is herein. All log and result files of the searches were exported for future reference and are available on our GitHub repository.

### FDR assessment by entrapment

We validated the theoretical FDR from the target-decoy approach during chromatogram library building by performing an entrapment experiment with *Pyrococcus furiosus*. In short, this is a way to additionally validate the target-decoy FDR estimation ^[13]^.Only peptides between 400 - 1000 *m/z* were considered and each source library requires a different *P. furiosus* input:

- A public *P. furiosus* dataset acquired on an LTQ-Orbitrap Velos (Thermo Fisher Scientific, Massachusetts, US) was used to supplement the Pan-Human DDA library (ProteomeXchange with identifier PXD001077)^[13]^. Database searching was performed on the resulting .mgf file with Mascot Daemon (version 2.6.1) using following search parameters: a maximum of one missed cleavage, peptide charges 2+ to 4+, peptide mass tolerance of 10 ppm, fragment ion tolerance of 0.5 Da, carbamidomethylation of Cysteine as fixed modification and oxidation of Methionine as variable modification. The resulting. DAT file was parsed into a. BLIB using the Skyline built-in tool BiblioSpec. The. BLIB file was parsed by EncyclopeDIA into a .dlib file. Finally, the resulting .dlib file (5.5k unique precursors) was combined with the already existing Pan-Human .dlib file of 194k peptides using EncyclopeDIA.
- For the FASTA database, we concatenated our FASTA with all 2052 *P. furiosus* UniProt entries (downloaded on June 13, 2018). Walnut parameters for library-free searching were set as described above, meaning that only 2+ and 3+ peptides without any variable modifications were considered. This translates into 168k *P. furiosus* precursors.
- For the predicted library, we converted this FASTA into a predicted *P. furiosus* spectral library using the MS²PIP Orbitrap-HCD model and our Elude RT model. Every 2+ and 3+ tryptic peptide in the proteome was predicted, with carbamidomethylation of Cysteine, and oxidation of Methionine set as respectively fixed and variable modifications. The *P. furiosus*. msp (224k precursors) was concatenated to the Human predicted .msp in EncyclopeDIA.

As decoys are generated by EncyclopeDIA, these were also appended for the *P. furiosus* proteins. All three source libraries were employed for searching the narrow window DIA data, i.e. to create a DIA-based chromatogram library. The *P. furiosus* fraction of the libraries was 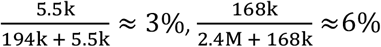 and 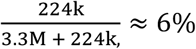 respectively. To account for this differential decoy fraction, the number of *P. furiosus* detections is multiplied by the inverse of their weights, using the following formula:

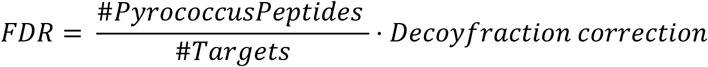

This corresponds to 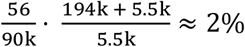 for the DDA source library, 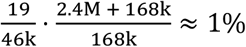 for the FASTA source library and 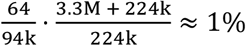 for the predicted source library. Note that the number of detected peptides (#targets) is slightly lower than the chromatogram libraries created without *P. furiosus* peptides (see main text). This corroborates the fact that increasing the number of false targets increases the statistical burden and thus number of false negatives, reducing the sensitivity of detection.

In the manuscript we claim the applicability of other deep learning predictors (e.g. DeepMass, Prosit) as an alternative to MS²PIP predicted libraries. To validate this claim we cloned the publicly available Pan-Human library using the Prosit webtool which is available from https://www.proteomicsdb.org/prosit/. Peptides containing more than 30 amino acids or with a charge state higher than 7 were manually removed from the list as this is required by Prosit. Normalized collision energy (NCE) was assumed to be 33 for all peptides. A similar clone of the Pan-Human library was made with the MS²PIP webtool using the pre-trained HCD model. After MS² peak intensity prediction, measured iRT values were parsed into both predicted libraries to remove the effect of retention time. Afterwards, the narrow window HeLa DIA data was searched against all three source libraries (Pan-Human, Prosit Clone and MS²PIP clone) separately using the settings described earlier in paragraph *DIA data analysis: EncyclopeDIA*. The results of these three searches were exported as the DDA, MS²PIP and Prosit chromatogram library, respectively. Next, three wide window HeLa DIA runs were searched with the three chromatogram libraries separately using the same settings as earlier. Again, the results were exported for further processing. The source and chromatogram libraries were converted to an OpenSWATH tsv by EncyclopeDIA, as this simplified parsing of the data. In accordance with the DIAmond DIAgram we calculated PCCs for each narrow and wide window DIA experimental spectrum and its DDA, MS²PIP and Prosit source and chromatogram spectrum. Only peptides containing at least 5 transitions were considered and *y1* ions were omitted.

**Figure S1.**
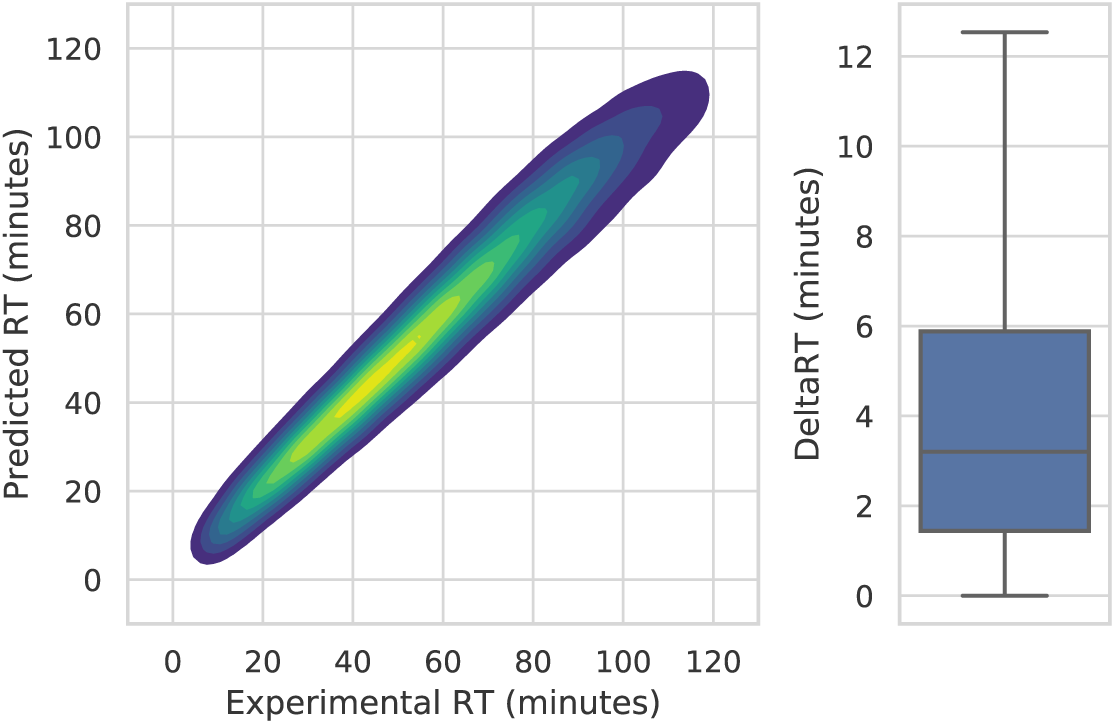
Evaluation of the trained Elude model. Left: Contour plot of all predicted and experimental retention times (RTs) in minutes. Right: Boxplot of all absolute differences between experimental and predicted RT (DeltaRT) in minutes. The box displays the first (Q1), second (Q2), and third (Q3) quartiles, the whiskers display Q1 - 1.5 times the interquartile range (IQR) and Q3 + 1.5 times the IQR, respectively. Outliers are not shown.

**Figure S2.**
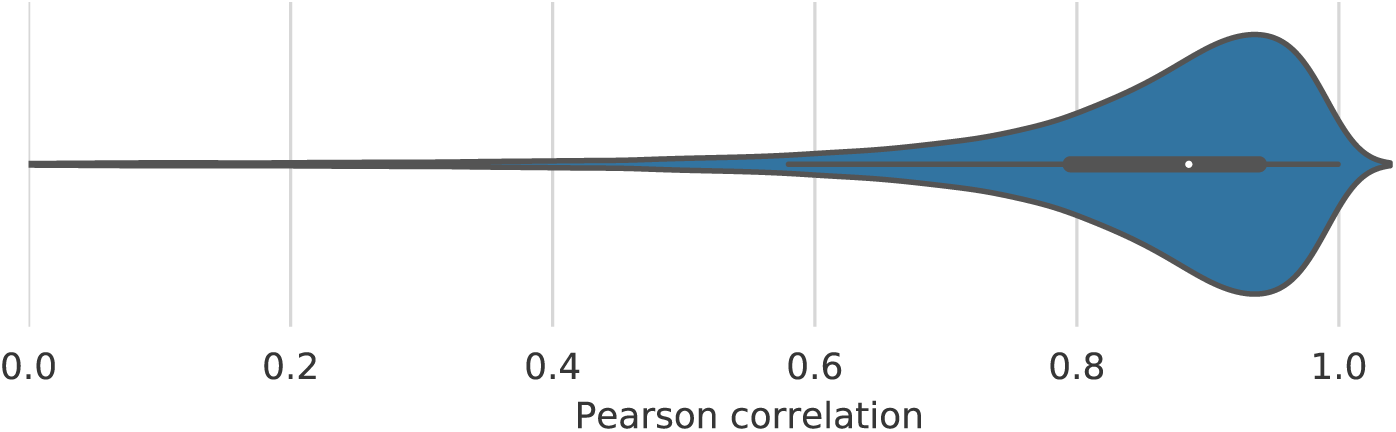
Pearson correlations between intensities of measured DDA and MS²PIP predicted fragments. Violin plot showing the distribution of Pearson correlation coefficients between the MS²PIP model predictions and the experimental spectra from the EncyclopeDIA article Hela DDA dataset.

**Figure S3:**
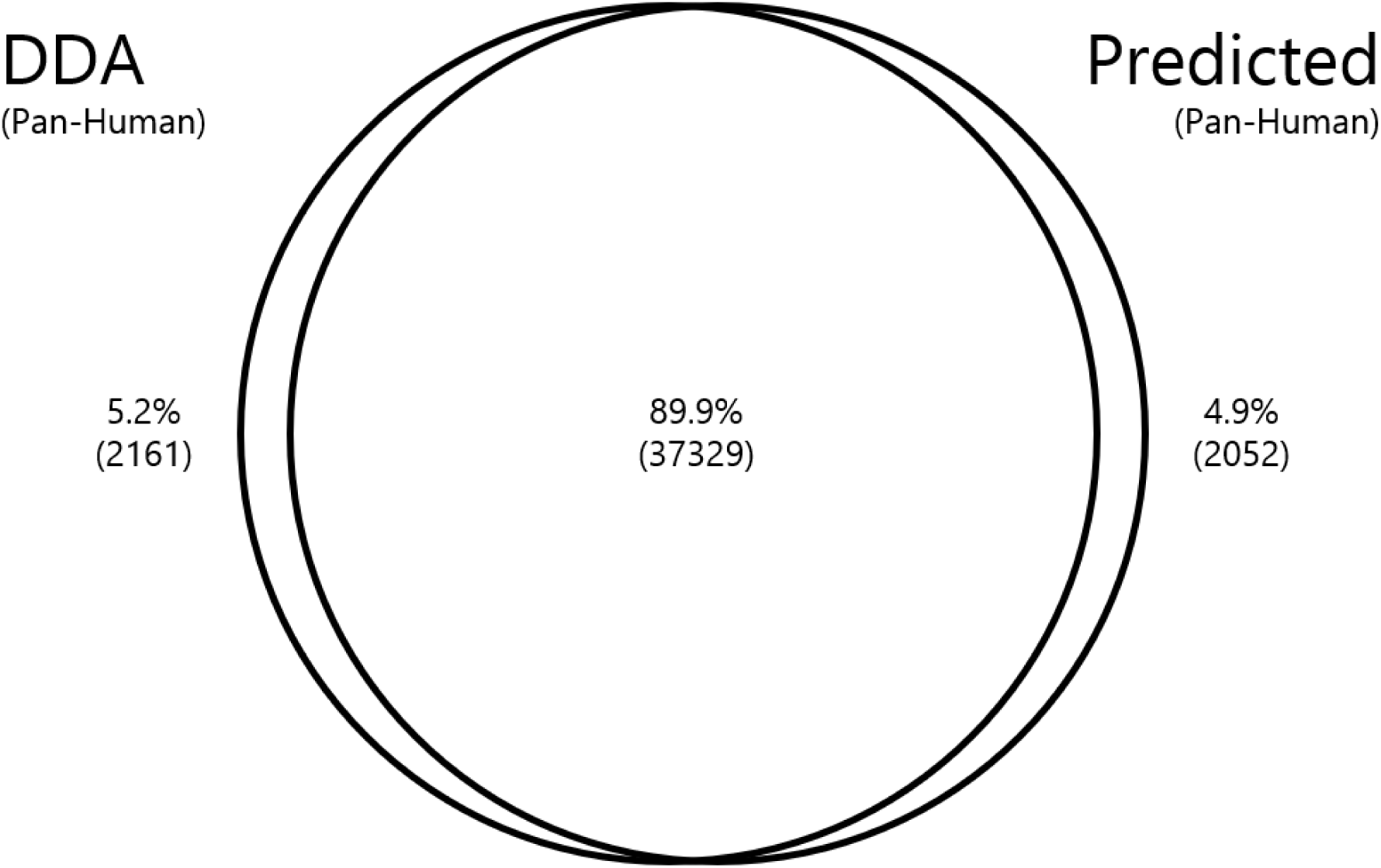
Overlap in peptides detected by DDA vs predicted chromatogram libraries. All peptides in a measured Pan-Human library were cloned by predicting their fragmentation spectra using MS2PIP and their retention times using ELUDE. A DIA library from a predicted library can extract peptides equally well from wide window DIA data compared to a DDA Pan-Human source library, a logical consequence of good quality predictions.

**Figure S4.**
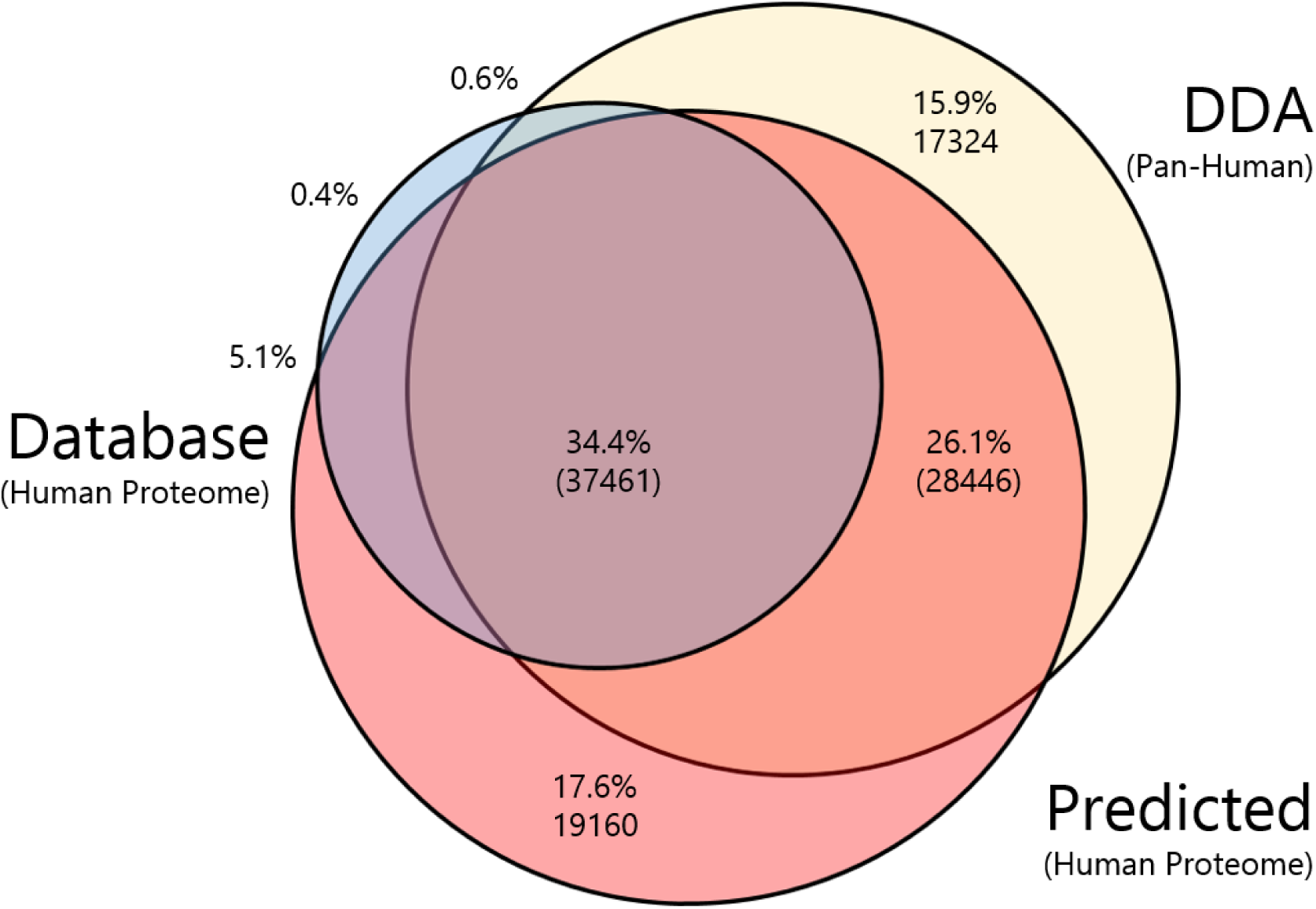
Overlap in peptide detection between all chromatogram libraries. A Venn-diagram showing the overlap in peptide sequence detections between the three DIA-based (DDA, Database and Predicted) chromatogram libraries.

**Figure S5.**
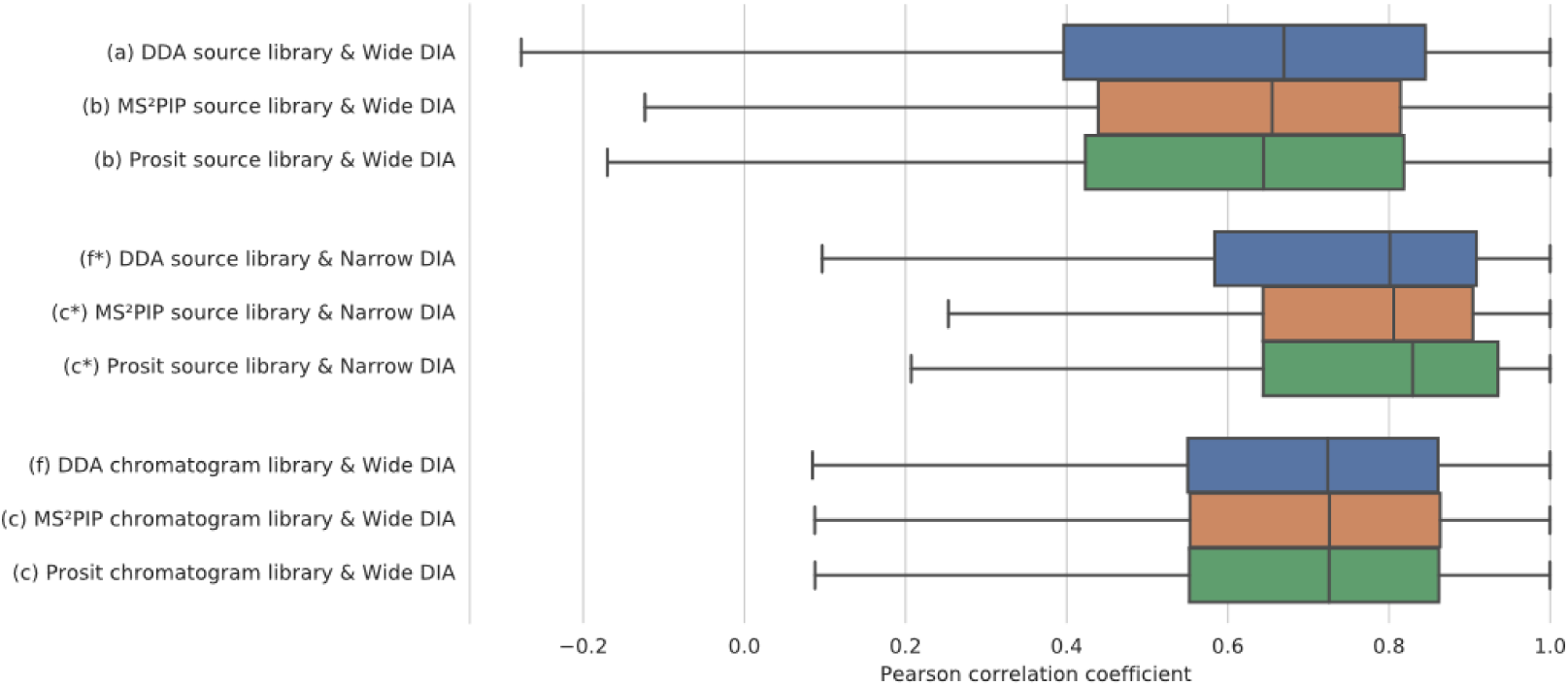
Boxplot showing the distribution of Pearson correlation coefficients between the experimental spectra from the Narrow and Wide-Window HeLa DIA data of the EncyclopeDIA article and the source libraries from DDA (a) or MS²PIP and Prosit (b), as well as the chromatogram libraries derived from DDA (f) or MS²PIP and Prosit (c). Letter annotations refer to the pathways in the DIAmond DIAgram (Figure 1). The overlapping boxplots of the three chromatogram libraries in the bottom clearly illustrate that calibration through narrow window DIA eliminates prior differences in (predicted) intensities.

## Notes

#### Summary of Updates

This version was resubmitted to include the minor revisions suggested by the reviewers.

https://github.com/brvpuyve/MS2PIP-for-DIA

